# Efficient CRISPR/Cas9 gene ablation in uncultured naïve mouse T cells for *in vivo* studies

**DOI:** 10.1101/730812

**Authors:** Simone Nüssing, Imran G. House, Conor J. Kearney, Stephin J. Vervoort, Paul A. Beavis, Jane Oliaro, Ricky W. Johnstone, Joseph A. Trapani, Ian A. Parish

**Author notes:** Correspondence: Ian Parish.

## Abstract

CRISPR/Cas9 technologies have revolutionised our understanding of gene function in complex biological settings, including T cell immunology. Current CRISPR-mediated gene deletion strategies in T cells require *in vitro* stimulation or culture that can both preclude studies of gene function within unmanipulated naïve T cells and can alter subsequent differentiation. Here we demonstrate highly efficient gene deletion within uncultured primary naïve murine CD8^+^ T cells by electroporation of recombinant Cas9/sgRNA ribonucleoprotein immediately prior to *in vivo* adoptive transfer. Using this approach, we generated single and double gene knock-out cells within multiple mouse infection models. Strikingly, gene deletion occurred even when the transferred cells were left in a naïve state, suggesting that gene deletion occurs independent of T cell activation. This protocol thus expands CRISPR-based probing of gene function beyond models of robust T cell activation, to encompass both naïve T cell homeostasis and models of weak activation, such as tolerance and tumour models.

## Introduction

Delineating the molecular mechanisms that underpin cellular pathways or processes is a major goal of most biological studies, and is typically achieved by probing the function of gene-deficient cells. In the field of CD8^+^ T cell biology, many major discoveries in areas ranging from homeostasis to effector cell differentiation, tolerance and exhaustion have been made using gene-deficient cells. However, generating such cells is often a bottleneck in the progress of CD8^+^ T cell research. Traditional approaches for the generation of gene-deficient cells typically involve whole genome or conditional knockout (KO) mouse models. While this remains the gold standard, it often requires complex mouse crosses that are slow and laborious, particularly when multiple genes are targeted. Cells derived from KO mice are also often not suitable for adoptive transfer studies because of cell rejection due to minor histocompatibility mismatches generated as a result of genetic drift or linked genetic loci carried on from the founder strain (1).

CRISPR (clustered, regularly interspaced short palindromic repeats)/Cas9 (CRISPR associated protein 9)-based gene deletion techniques have recently revolutionised our ability to rapidly generate gene deficient cells (2). In this system, guide RNAs, or more recently single guide RNAs (sgRNAs), are used to target the Cas9 nuclease to a genomic region of interest, leading to double-stranded DNA breaks that trigger disruptive insertions or deletions (3). Transduction of Cas9-expressing cells with viral delivery vectors encoding CRISPR guide RNAs is an efficient approach for triggering CRISPR/Cas9-mediated gene deletion within a target cell population (4-6). While this strategy has worked well for T cells (5), it has a number of drawbacks that limit its utility. First, transduction with viral vectors typically requires *in vitro* T cell activation and culture, which both alters *in vivo* differentiation (particularly settings involving weak activation, such as tumour and tolerance models) and precludes the use of these technologies for the study of naïve T cell homeostasis. While recently reported bone marrow chimera approaches can address some of these problems (7), this strategy remains time consuming and unsuitable for studying genes required for thymic T cell development. Second, persistent Cas9 expression both increases the likelihood of off-target effects (8, 9), and is not suitable for mouse *in vivo* adoptive transfer approaches, as the immunogenic Cas9 protein can cause cell rejection (10, 11).

To overcome these problems, electroporation of cells with recombinant Cas9/sgRNA ribonucleoprotein (RNP) has recently been described (12-14), with the use of chemically modified guides leading to high editing efficacy (14). This strategy leads to highly efficient gene deletion, and avoids the problems associated with persistent Cas9 expression. Moreover, a recent study demonstrated that this approach works well *in vitro* in both naïve and activated mouse and human T cells (15). The capacity to delete genes within naïve T cells was particularly significant, as it implied that this approach could avoid the caveats associated with *in vitro* T cell activation in other systems. Nevertheless, in this study, resting murine T cells still had to be either activated via αCD3/αCD28 shortly after electroporation, or pre-cultured in IL-7, for efficient gene deletion (15). It remains unknown whether this approach results in efficient gene deletion within *in vivo* mouse T cell adoptive transfer systems, where activation signals are not as rapidly delivered or as strong as *in vitro* α-CD3/α-CD28 treatment. Moreover, while the IL-7 culture step did not lead to overt T cell activation (15), it does raise questions about the suitability of this approach within *in vivo* experimental models, such as exhaustion models, where exogenous IL-7 can alter differentiation (16, 17).

Here we demonstrate highly efficient CRISPR/Cas9-mediated gene deletion within *in vivo* models by sgRNA/Cas9 RNP electroporation into primary, uncultured and unstimulated naïve TCR-transgenic CD8^+^ T cells prior to adoptive transfer. We validate this model using multiple genes and multiple TCR transgenic T cell specificities, in two well-established infection models: chronic Lymphocytic choriomeningitis virus Clone 13 (LCMV-Cl13) and ovalbumin-transgenic *Listeria monocytogenes* (LM-OVA). Adoptively transferred cells underwent clonal expansion, lost target protein expression at high efficiency, and were readily detectable *in vivo* after infection. The high efficiency of this approach also enabled efficient generation of cells in which two genes were disrupted. Strikingly, gene deletion was equally efficient when naïve T cells were injected into recipients and “parked” without stimulation, implying that gene deletion occurs independently of T cell activation. Collectively, these findings validate naïve T cell electroporation for *in vivo* studies, and enable rapid probing of gene function without the caveats of *in vitro* cell culture. This method additionally facilitates rapid *in vivo* CRISPR-based studies on naïve T cell homeostasis and weak T cell activation models.

## Materials and Methods

### Mice

6-10 week old male or female mice were used for all described experiments, with mice housed in the Peter MacCallum Cancer Centre Animal Core Facility under specific pathogen-free conditions. CD45.2^+^ C57BL/6 (B6) recipient mice were obtained from the Walter and Eliza Hall Institute Kew Animal Facility (Victoria, Australia), while congenic CD45.1^+^ TCR-transgenic P14 (18) and CD45.1^+^ TCR-transgenic OT-I (19) donor mice were bred in-house. All animal work was in accordance with protocols approved by the Peter MacCallum Cancer Centre Animal Experimentation Ethics Committee (Protocol E597), and current guidelines from the Australian Code of Practice for the Care and Use of Animals for Scientific Purposes.

### Naïve CD8^+^ T cell enrichment

Single cell suspensions were prepared from spleen and pooled lymph nodes (LNs) isolated from either CD45.1^+^ congenic TCR-transgenic P14 or OT-I donor mice, and CD8^+^ T cells were then magnetically enriched using the EasySep™ Mouse CD8^+^ T Cell Isolation Kit (STEMCELL™ Technologies) according to manufacturer’s instructions. Purity of enriched naïve CD8^+^ T cells was determined by flow cytometry staining with anti-mouse CD8α-PerCP (Clone 53-6.7; Biolegend, CA, USA), Vα2-PE (Clone B20.1), Ly5.1-FITC (Clone A20) (both BD Biosciences, NJ, USA), with samples acquired on a BD FACSCanto™ II (BD Biosciences).

### CRISPR/Cas9 electroporation, adoptive transfer, and mouse infection

sgRNAs targeting the murine *Cd3e* (AGGGCACGUCAACUCUACAC), *Thy1* (encoding CD90) (CCUUGGUGUUAUUCUCAUGG) or *Pdcd1* (GACACACGGCGCAAUGACAG) genes, and the mouse genome non-targeting *Ctrl* sgRNA (GCACUACCAGAGCUAACUCA), were purchased from Synthego (CRISPRevolution sgRNA EZ Kit, Synthego). For sgRNA/Cas9 RNP formation, 1 µl of sgRNA (0.3 nmol/µl in nuclease-free H_2_O) was incubated with 0.6µl of Alt-R^®^ S.p. Cas9 Nuclease V3 (10 mg/ml in 50% glycerol, Integrated DNA Technologies) in nuclease-free H_2_O in a final volume of 5 µl for 10 min at room temperature. In double KO experiments, 1 µl of each sgRNA (2µl total) and 0.6µl Cas9 were used, with the final volume also adjusted to 5 µl as above. After washing in PBS, 2-10×10^6^ *ex vivo* enriched CD8^+^ T cells per electroporation condition were pelleted, resuspended in 20µl P3 buffer (P3 primary cell 4D-Nucleofector™ X Kit S, Lonza Ltd), and mixed with the pre-prepared 5 µl sgRNA/Cas9 RNP prior to immediate electroporation using the P3 primary cell 4D-Nucleofector™ X Kit S electroporation wells and Lonza 4D-Nucleofector™ System (Pulse DN 100). 130 µl RPMI (Gibco) containing 10% Fetal Calf Serum, glutamine and antibiotics were added to cells in electroporation wells, and cells were rested for 10 min at 37°C and 5% CO_2_. T cells were then counted and adoptively transferred intravenously into B6 recipient mice, with 5×10^3^ P14 T cells transferred per recipient mouse for LCMV-Cl13 infection, 5×10^4^ OT-I cells per recipient mouse for LM-OVA infection, and 1-2×10^6^ cells per mouse for naïve cell “parking” experiments. For infections, mice were intravenously injected with either 2.5x×10^6^ p.f.u. LCMV-Cl13 or 5×10^4^ c.f.u. LM-OVA.

### Flow Cytometric Analysis

For flow cytometric analysis, single cell splenic suspensions were stained on ice in PBS containing 2.5% Fetal Calf Serum and 0.1% sodium azide using the following anti-mouse antibodies and reagents: Fixable Viability Stain 620, CD8-BUV395 (Clone 53-6.7), CD45.1-FITC (Clone A20), CD3-APC (Clone 145-2C11), Thy1.2-PE (Clone 53-2.1) (all from BD Biosciences) and PD-1-BV785 (Clone 29F.1A12; BioLegend). Samples were acquired on a BD LSRFortessa X-20 (BD Biosciences) and analysed using FlowJo software (TreeStar) and GraphPad Prism (GraphPad Software). All *p* values were calculated using an unpaired One-way ANOVA with a Tukey post-test, except Fig. 4C where a two-tailed unpaired student’s T test was used.

**FIGURE 1.**
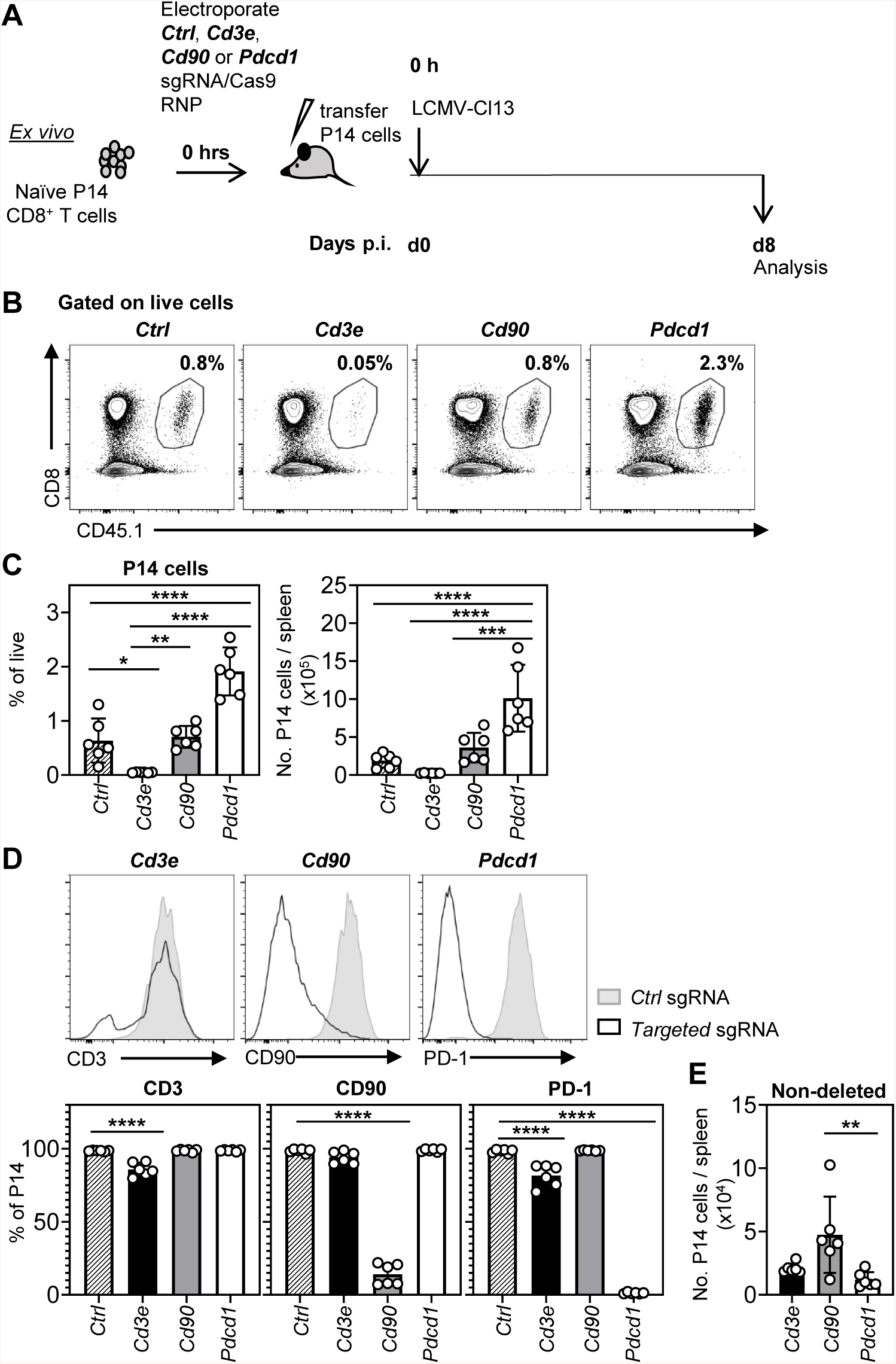
Efficient CRISPR/Cas9-mediated gene deletion in adoptively transferred naïve P14 T cells after LCMV-Cl13 infection. (**A**) Purified naïve CD8^+^ CD45.1^+^ P14 cells were electroporated with targeting or non-targeting sgRNA/Cas9 RNPs and 5×10^3^ P14 cells were immediately transferred into congenic CD45.2^+^ B6 recipient mice (0 h). Recipient mice were simultaneously infected with 2.5×10^6^ p.f.u. of LCMV-Cl13. Gene deletion efficiency and T cell expansion within splenic P14 cells were analysed on day 8 p.i. (**B**) Representative profiles and (**C**) pooled percentages (left) and numbers (right) of transferred CD45.1^+^ P14 CD8^+^ T cells electroporated either with non-targeting *Ctrl, Cd3e-, Cd90-* or *Pdcd1*-targeting sgRNA/Cas9 RNPs. (**D**) Representative histograms (top) and pooled percentages (bottom) of CD3, CD90 and PD-1 surface expression in sgRNA/Cas9 electroporated CD45.1^+^ P14 CD8^+^ T cells. (**E**) Total number of CD45.1^+^ P14 CD8^+^ T cells that retained expression of the respective targeted surface molecule on day 8 p.i. (“non-deleted”). Data and representative FACS plots are from two independent experiments with a total of n=6 mice per group. \**p* < 0.05, \*\**p* < 0.01, \*\*\*\**p* < 0.0001. Bars depict mean, while error bars represent SD.

**FIGURE 2.**
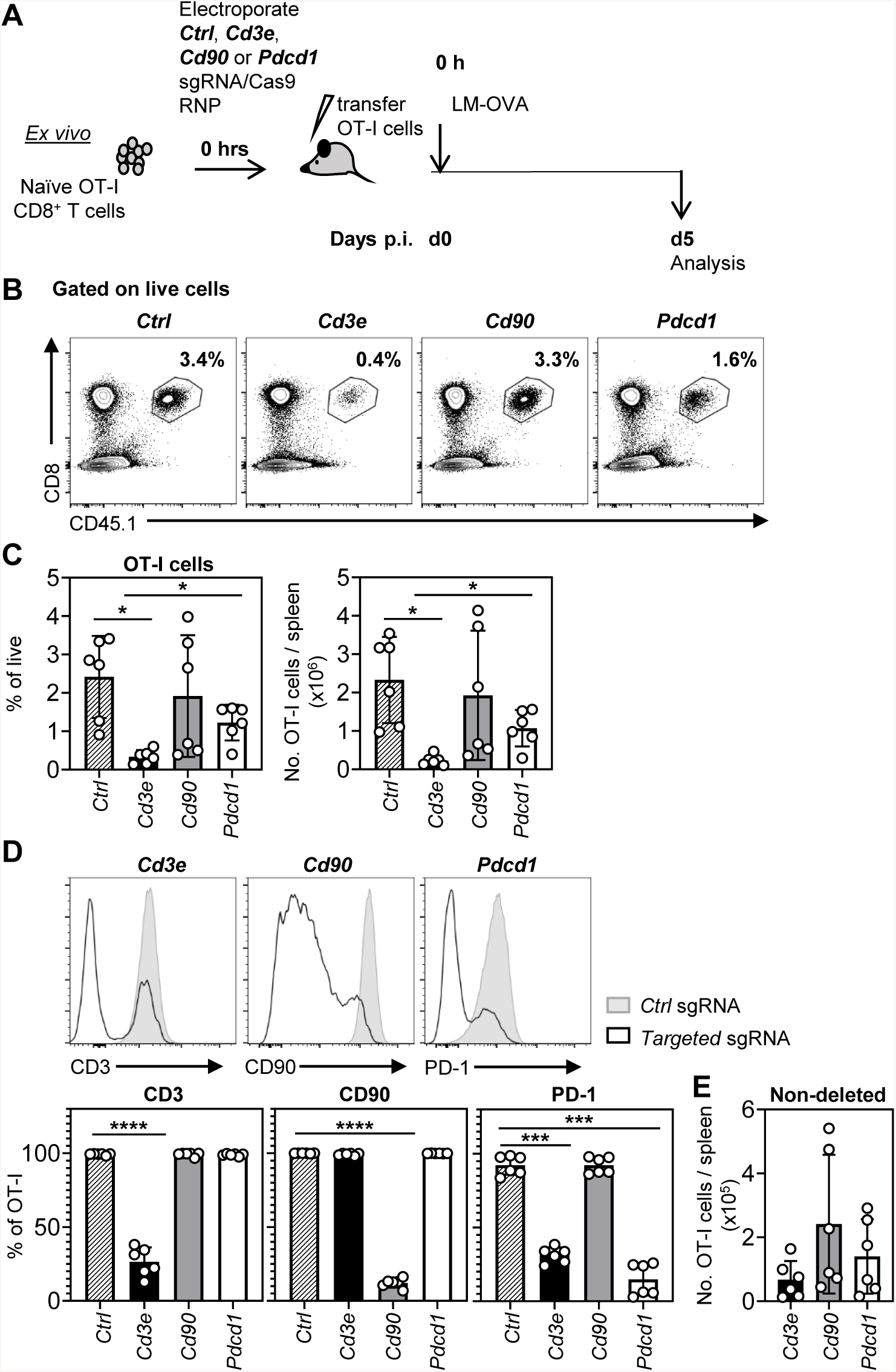
Efficient CRISPR-mediated gene loss in adoptively transferred naïve OT-I T cells after LM-OVA infection. (**A**) Purified naïve CD8^+^ CD45.1^+^ OT-I cells were electroporated with targeting or non-targeting sgRNA/Cas9 RNPs and 5×10^4^ OT-I cells transferred into congenic CD45.2^+^ B6 recipient mice that were simultaneously infected with 5×10^4^ c.f.u. of LM-OVA. Gene deletion efficiency and T cell expansion was analysed on day 5 p.i.. (**B**) Representative profiles and (**C**) pooled percentages (left) and numbers (right) of transferred CD45.1^+^ OT-I CD8^+^ T cells electroporated with the sgRNAs used in Fig. 1B. (**D**) Representative histograms (top) and pooled percentages (bottom) of CD3, CD90 and PD-1 surface expression in sgRNA/Cas9 electroporated CD45.1^+^ OT-I CD8^+^ T cells. (**E**) Total number of “non-deleted” CD45.1^+^ OT-I CD8^+^ T cells as defined in Fig. 1E. Data and representative FACS plots are from two independent experiments with a total of n=6 mice per group. *p < 0.05, \*\*\**p* < 0.001, \*\*\*\**p* < 0.0001. Bars depict mean, while error bars represent SD.

**FIGURE 3.**
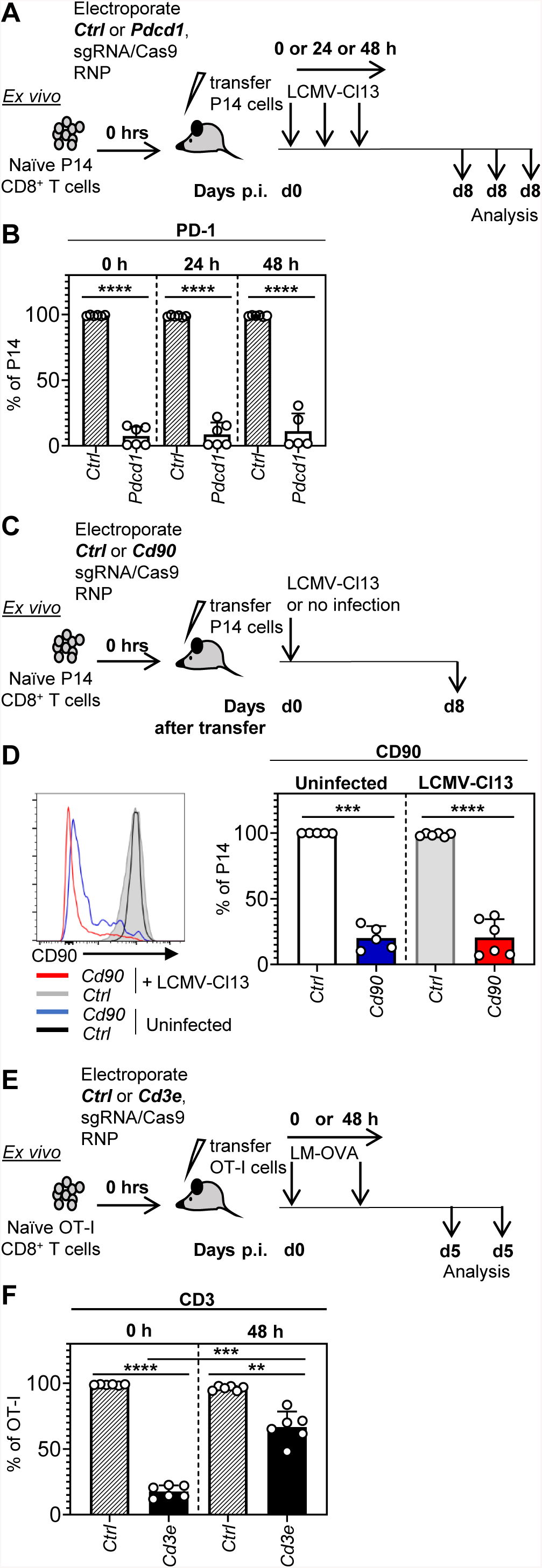
CRISPR/Cas9-mediated gene deletion occurs within transferred naïve CD8^+^ T cells independent of infection. (**A and B**) The experiment in Fig. 1A was repeated using *Ctrl* or *Pdcd1*-targeting sgRNA/Cas9 RNPs, except that LCMV-Cl13 infection was either immediate or delayed until 24 h or 48 h after transfer (**A**). (**B**) illustrates PD-1 levels in the transferred P14 cells at day 8 p.i. (**C and D**) Naïve P14 cells were electroporated with either *Ctrl*- or *Cd90*-sgRNA/Cas9 RNPs, and transferred into recipient B6 mice that were either immediately LCMV-Cl13 infected (transfer of 5×10^3^ cells) or left uninfected (transfer= of 1-2×10^6^ cells) (**C**). (**D**) shows representative (left) and pooled (right) CD90 deletion efficiency on day 8 p.i. (**E and F**) The experiment in Fig. 2A was repeated using non-targeting *Ctrl* or *Cd3e*-targeting sgRNA/Cas9 RNPs, except that LM-OVA infection was either immediate or delayed 48 h after transfer (**E**). (**F**) shows CD3 expression at day 5 p.i.. Data and representative FACS plots are from two independent experiments with a total of 421 n=5-6 mice per group. \*\**p* < 0.01, \*\*\**p* < 0.001, \*\*\*\**p* < 0.0001. Bars depict mean, while error bars represent SD.

**FIGURE 4.**
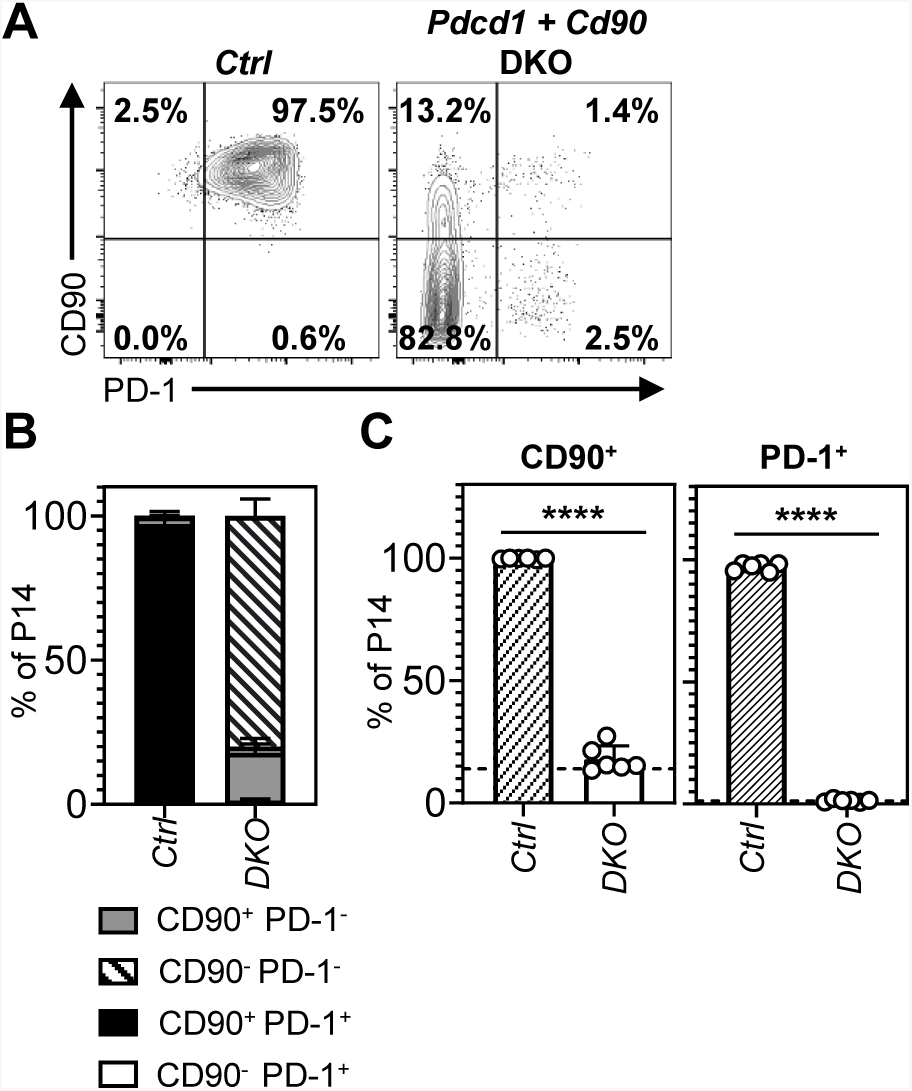
Efficient CRISPR/Cas9-mediated generation of double KO CD8^+^ T cells. The experiment outlined in Fig. 1A was repeated, except that P14 cells were electroporated with either non-targeting *Ctrl* or combined *Pdcd1*- and *Cd90*-targeting sgRNA/Cas9 RNPs. Representative (**A**) and pooled (**B and C**) proportions of cells lacking PD-1 and/or CD90 are shown. (**B**) shows proportion of cells from each quadrant in the plots illustrated in (**A**). (**C**) illustrates the proportion of cells expressing CD90 (regardless of PD-1 expression) or PD-1 (regardless of CD90 expression). Dotted lines in the CD90^+^ and PD-1^+^ plots indicate the knock-out efficiency seen in cells given each guide singly in Fig. 1D. Data and representative FACS plots are from two independent experiments with n=6 mice per group. \*\*\*\**p* < 0.0001. Bars depict mean, while error bars represent SD.

## Results

### Effective gene knockout in CRISPR-edited adoptively transferred naïve CD8^+^ P14 T cells after LCMV-Cl13 infection

Previous work has suggested that sgRNA/Cas9 RNP electroporation into cultured naïve T cells can result in efficient gene deletion (15). We sought to modify this strategy to both test its utility within *in vivo* mouse models, and also avoid *in vitro* culture. To this end, naïve CD8^+^ LCMV-specific CD45.1^+^ P14 cells were purified from donor mice and immediately electroporated without any *in vitro* activation or cytokine culture step. To further increase the accessibility and efficiency of this approach, we used readily available, commercially synthesized, chemically modified sgRNAs that are specifically optimised for electroporation, and a commercially available recombinant *Streptococcus pyogenes* (S.p.) Cas9 nuclease. The electroporated P14 cells were then immediately transferred into recipient CD45.2^+^ B6 mice that were simultaneously infected with chronic LCMV-Cl13. At day 8 post-infection (p.i.) we measured splenic P14 cell expansion and gene deletion by flow cytometry (Fig. 1A). For these proof-of-concept experiments, we chose 3 different classes of gene to target: a gene required for TCR signalling and expansion (*Cd3e*), one that restrains CD8^+^ T cell expansion in this model (*Pdcd1*) and another whose deletion is expected to have no effect on expansion (*Cd90*; *Thy1* gene). Deletion of these genes results in predictable biological effects that CRISPR-mediated deletion should phenocopy, and all protein products of these genes can be easily traced with validated antibodies by flow cytometry.

Consistent with the known phenotypes associated with gene deficiency, relative to P14 cells treated with a control non-targeting (*Ctrl*) sgRNA, *Cd3e* sgRNA treated cells had a severe expansion deficiency, *Cd90* sgRNA treated cells expanded normally, while *Pdcd1* sgRNA treated cells exhibited augmented expansion (Fig. 1B and C). This was associated with high deletion efficiency within *Cd90* and *Pdcd1* targeted cells relative to control cells (86% and 99% KO cells respectively) (Fig. 1D). Notably, only 14.2% of *Cd3e* targeted P14 cells had lost CD3 protein expression, however this is likely because the cells that failed to delete *Cd3e* preferentially expanded. Consistent with this idea, the total number of P14 T cells that failed to undergo gene deletion (“non-deleted”) did not significantly differ between *Cd3e*, and *Cd90* or *Pdcd1*, targeted P14 cells (Fig. 1E). Non-deleted cell number differed significantly between *Cd90* and *Pdcd1* targeted cells, although this likely reflected the lower gene editing efficiency of the *Cd90* sgRNA. Thus, uncultured sgRNA/Cas9 RNP electroporated naïve P14 cells efficiently delete genes when activated by LCMV-Cl13 *in vivo*.

### High efficiency gene deletion within a different target T cell population and infection model

To test if high gene deletion efficiency could be observed within different target T cell populations and in a different infection model, we performed similar experiments using ovalbumin-specific CD8^+^ CD45.1^+^ OT-I TCR-transgenic T cells adoptively transferred into B6 mice simultaneously infected with LM-OVA. As PD-1 expression is only evident early during infection in this model (20), splenic OT-I cells were analysed at day 5 p.i. (Fig. 2A). Again, *Cd3e* targeted cells exhibited an expansion deficiency, whereas *Cd90* targeted cells expanded normally (Fig. 2B and C). *Pdcd1* targeted cells did not exhibit significantly altered expansion, consistent with previous reports describing either a neutral or positive role for the PD-1/PD-L1 axis in OT-I expansion within this model (21, 22). Again, high deletion efficiency was observed within *Cd90* and *Pdcd1* targeted cells (88% and 85% respectively) (Fig. 2D). Similar to Fig. 1E, comparable numbers of “non-deleted” cells were again observed in all conditions (Fig. 2E), however a much higher proportion (74%) of gene deficient *Cd3e-*sgRNA electroporated OT-I T cells were recovered (Fig. 2D). As CD3-deficient cells should not respond to antigen and expand, we speculated that the increased proportion of CD3-deficient cells was due to the earlier (day 5) time-point in these experiments; it is likely that there is a lag between initial gene deletion and protein loss, meaning KO cells can initiate expansion prior to eventual protein loss. Nevertheless, expansion was still impaired within this group, and overall, these data illustrate that this experimental approach is similarly efficient in an independent *in vivo* setting.

### Comparable gene deletion in naïve P14 T cells transferred into uninfected mice

Previous *in vitro* work has suggested that immediate strong T cell activation is required for efficient gene deletion within electroporated, uncultured naïve T cells (15). As Cas9 protein is likely rapidly lost from cells after electroporation, we sought to define how long infection could be delayed post-transfer without compromising deletion efficiency. To this end, we undertook a time-course study in which B6 mice given *Ctrl* or *Pdcd1* sgRNA electroporated CD45.1^+^ P14 cells were infected with LCMV-Cl13 either immediately (0 h), 24 h, or 48 h after P14 transfer, with PD-1 expression analysed at day 8 p.i. (Fig. 3A). Surprisingly, there was no significant difference in gene deletion efficiency regardless of how long infection was delayed (Fig. 3B). This could be explained by either sgRNA/Cas9 RNP persistence in P14 cells, or alternatively because gene editing may occur in naïve P14 cells independent of T cell activation. To discriminate between these possibilities, we examined whether deletion of a gene constitutively expressed in naïve T cells (*Cd90*) would still occur within adoptively transferred naïve P14 cells that were kept in a naïve state. *Ctrl* or *Cd90* targeted CD45.1^+^ P14 cells were either transferred into B6 mice immediately infected with LCMV-Cl13, or B6 mice that were left uninfected (Fig. 3C). Strikingly, when we measured CD90 expression at day 8 post-transfer, no differences were observed in the proportion of CD90 deficient P14 cells recovered from infected or uninfected recipients (79.5% vs 79.9% respectively; Fig. 3D). Thus, this approach can be used for efficient gene deletion within naïve CD8^+^ T cells without any prior *in vitro* culture or conditioning.

The capacity of CRISPR/Cas9 gene editing to occur within unmanipulated naïve CD8^+^ T cells in this system meant that delaying infection may be beneficial, as it would enable protein depletion prior to activation. We previously observed a large proportion of expanded CD3 KO OT-I cells in LM-OVA infection (Fig. 2D), and we speculated that this KO population would be reduced if infection was delayed to enable protein depletion prior to activation. Consistent with this idea, when we repeated this experiment but delayed infection until 48 h after OT-I transfer (Fig. 3E), a much lower proportion of CD3 KO cells were recovered (Fig. 3F). Thus, gene deletion within naïve T cells via this method can be employed to ensure protein depletion prior to activation.

### CRISPR-mediated deletion of multiple genes within transferred naïve CD8^+^ T cells

Given the high efficiency of gene deletion using this approach, we next attempted to generate double KO cells by combined electroporation of two guides (targeting *Cd90* and *Pdcd1*) into CD45.1^+^ P14 T cells prior to transfer and LCMV-Cl13 infection. A very high proportion (80%) of cells lacked both proteins (Fig. 4A and B) and, importantly, the knockout efficiency of each individual sgRNA was comparable to when the guides were introduced alone in previous experiments (dashed lines) (Fig. 4C). This indicated that there was no competitive inhibition between sgRNAs simultaneously introduced into the same cell, and validated that this experimental approach can simultaneously target multiple genes.

## Discussion

A major limitation of current CRISPR/Cas9-mediated gene editing protocols in T cells has been the requirement for *in vitro* culture, which restricts the experimental questions that can be addressed. Here we were able to adapt a previously described CRISPR-based gene deletion strategy in murine T cells (15) to achieve high efficiency gene deletion in adoptively transferred naïve CD8^+^ T cells *in vivo*. Due to high knockout efficiency, our described approach did not require selection or sorting of targeted cells, and, importantly, no prior naïve T cell culture was required for gene deletion. This approach provides a rapid CRISPR-based deletion strategy for use in *in vivo* naïve T cell homeostasis studies, as well as *in vivo* adoptive transfer models (e.g. peripheral self-tolerance models) that are compromised by the *in vitro* culture and activation steps required in other CRISPR protocols. This is also, to our knowledge, the first report describing the use of naïve T cell electroporation for gene deletion within *in vivo* adoptive transfer mouse models.

Our most striking finding was that our approach leads to highly efficient gene deletion within uncultured naïve T cells *in vivo*. Previous CRISPR/Cas9 gene deletion approaches in T cells required either T cell activation, or culture with supraphysiological levels of IL-7, to facilitate either retroviral transduction (6), plasmid electroporation (23) or efficient gene deletion after sgRNA/Cas9 RNP electroporation (15). These culture steps make these approaches unsuitable for certain adoptive transfer models, where prior activation or cytokine culture could alter *in vivo* T cell differentiation (eg. tolerance and tumour models). In contrast, we conducted all of our experiments without prior *in vitro* activation or conditioning of the target naïve T cells. We find that when CRISPR/Cas9 electroporated T cells are used within *in vivo* mouse adoptive transfer models, not only is activation by *in vivo* infection sufficiently strong and rapid to facilitate gene deletion, but naïve cells transferred *in vivo* exhibit strong gene deletion. This contrasts with previous findings that naïve T cells cultured without activation do not delete genes as efficiently as that seen after T cell activation (15). This discrepancy appears to be in part due to differences between adoptively transferred versus IL-7-stimulated naïve T cells; we have preliminary data that naïve P14 cells transferred into mice exhibited more efficient *Cd90* gene deletion than the same cells cultured in IL-7 (data not shown). Regardless of mechanism, this finding provides a solution to the complications and biases introduced into experiments by *in vitro* culture steps prior to adoptive T cell transfer.

Additionally, we achieved efficient compound knockout of multiple genes using this method without competition between different gene targeting sgRNAs. This provides a far more rapid approach for generating compound KO cells than conventional mouse breeding strategies, and is superior to retroviral CRISPR-based compound KO strategies that are often inefficient and/or limited by vector size (4). Our observation that there is no competition between co-delivered CRISPR/Cas9 RNPs also implies that this strategy could be used to delete more than two genes, which is currently not easily achieved with retroviral-based strategies. Finally, our findings highlight that the half-life of the targeted protein needs to be considered when using these approaches. When working with a protein that has a long half-life, gene deletion within naïve T cells prior to activation should be strongly considered. Conversely, our approach may be useful to determine the *in vivo* half-life of the protein expressed by a targeted gene. Collectively, the method reported in this paper is thus a powerful new technique that enables the use of CRISPR-based approaches in models that were previously not amenable to these strategies.

## Acknowledgements

We thank the Peter MacCallum Cancer Centre Animal Core facility for breeding and maintenance of the mice, the Peter MacCallum Cancer Centre Flow Cytometry Facility, and the Peter MacCallum Cancer Centre Genotyping Laboratory for genotyping of mice.

## Footnotes

This study was supported by Human Frontier Science Program Young Investigators Grant RGY0065/2018, and Program, Fellowship and Project grant support from the National Health and Medical Research Council (NHMRC) of Australia.

## Abbreviations

CRISPR: clustered, regularly interspaced short palindromic repeats;
Cas9: CRISPR associated protein 9;
*Ctrl*: non-targeting control sgRNA;
sgRNA: single guide RNA;
LCMV-Cl13: Lymphocytic choriomeningitis virus Clone 13 strain;
LM-OVA: ovalbumin-transgenic *Listeria monocytogenes*

